# Generate a new crucian carp (*Carassius auratus*) strain without intermuscular bones by knocking out *bmp6*

**DOI:** 10.1101/2022.11.28.518130

**Authors:** Youyi Kuang, Xianhu Zheng, Dingchen Cao, Zhipeng Sun, Guangxiang Tong, Huan Xu, Ting Yan, Shizhan Tang, Zhongxiang Chen, Tingting Zhang, Tan Zhang, Le Dong, Xiaoxing Yang, Huijie Zhou, Weilun Guo, Xiaowen Sun

## Abstract

Elimination of intermuscular bones (IMBs) is vital to the aquaculture industry of cyprinids. In our previous study, we characterized *bmp6* as essential in the development of IMBs in zebrafish. Knockout of *bmp6* results in the absence of IMBs in zebrafish without affecting growth and reproduction. Therefore, we hypothesized that *bmp6* could be used to generate new cyprinid strains without IMBs by gene editing. In this study, we established a gene editing strategy for knocking out the two orthologs of *bmp6* in diploid crucian carp (*Carassius auratus*). We obtained an F_3_ population with both orthologs knocked out, in which no IMBs were detected by bone staining and X-ray, indicating that the new strain without IMBs (named WUCI strain) was successfully generated. Furthermore, we extensively evaluated the performance of the new strain in growth, reproduction, nutrient components, muscle texture and structure, and metabolites in muscle. The results showed that the WUCI strain grew faster than the wild-type crucian carp at 4-month-age. The reproductive performance and flesh quality did not show significant differences between the WUCI strain and wild-type crucian carp. Moreover, the metabolomics analysis suggested that the muscle tissues of the WUCI strain significantly enriched some metabolites belonging to the Thiamine metabolism, Nicotinate and Nicotinamide Metabolism pathway, which plays beneficial effects in anti-aging, anti-oxidant, and anti-radiation damage. In conclusion, we established a strategy to eliminate IMBs in crucian carp and obtain a WUCI strain whose performance was expected compared to the wild-type crucian carp; meanwhile, the WUCI strain enriched some beneficial metabolites to human health in muscle tissue. This study is the first report that a farmed cyprinid strain without IMBs, which could be stably inherited, was obtained worldwide; it provided excellent germplasm for the cyprinids aquaculture industry and a useful molecular tool for eliminating IMBs in other cyprinids.

## 1. Introduction

Cyprinids are the most important aquaculture species in China, with an annual production of about 27 million tons, occupying around 76% of total freshwater fish aquaculture production. However, the intermuscular bones (hereafter IMBs) severely cause food inconvenience and may even lead to physical injuries such as being stuck in the throat. The processing of fish products (such as fish balls) is also hindered. The genetic breeding of eliminating IMBs of cyprinids has become an increasing interest in China(Gui et al., 2021; Nie et al., 2020; Sun and Zhu, 2019).

The genetic breeding of strains without IMBs in cyprinids began in the 1960s (Meske, 1968), but there has been no progress until the last decade. With the development of biotechnology, some progress has been made in the genetic breeding of IMBs. For example, Xu et al. (2015) found a mutant with normal growth and absence of IMBs from a gynogenesis population of grass carp(*Ctenopharyngodon idellus*); Peruse et al.(2017) obtained a population with IMBs deficient in *Colossoma macropomum* by X-ray scan; Wu et al. (2020) obtained a population with a 5.7% reduction in the count of IMBs by crossing *Megalobrama amblycephala* and *Culter alburnus*, indicating that varieties with less count of IMBs can be obtained by crossbreeding. Lin et al.(2013) reported that hybrid crucian carp (*Carassius auratus*) had a smaller number of IMBs than wild crucian carp, and the count of IMBs of crucian carp could be reduced by polyploid breeding and crossbreeding methods; Cao et al. (2015) concluded that mirror carp (*Cyprinus carpio*) could be an excellent candidate for genetic selection to reduce IMBs. In terms of genetic and molecular breeding, Xiong et al. (2019) found that the heritability of the number of IMBs differed in different parts of blunt snout bream (*M. amblycephala*); Wan et al. (2019) identified single nucleotide polymorphism (SNP) markers associated with the count of IMBs in *M. amblycephala*. The counts of IMBs were significantly different among common carp (*C. carpio*) species/strains and families (Cao et al., 2015; Tang et al., 2020). However, the heritability of IMBs was low, with a value of 0.15 (Tang et al., 2020). Eight significant QTL intervals were obtained in the QTL analysis of the number of IMBs in mirror carp, which explained 7.7-25.9% of the phenotypic variation(Tang et al., 2020).

From the above statements, it is clear that genetic improvement of IMBs can be carried out by artificial selection, cross breeding, gynogenesis, polyploid breeding, and molecular marker-assisted selection; however, there is still no breakthrough in reducing more than 50% of IMBs. With the development of gene editing technology, the genes and regulatory elements affecting the development of IMBs have been characterized. For example, Nie et al. (2021) knocked out *scxa* in zebrafish and obtained a mutant with more than 70% reduction in the count of IMBs, rib hypoplasia, and defective septal development(Kague et al., 2019; Nie et al., 2021). In our previous study, we characterized that *bmp6* regulates the development of IMBs in zebrafish(Xu et al., 2022), and its knockout causes the complete absence of IMBs in zebrafish without affecting the growth, muscle, and bone development of zebrafish (Jian et al., 2019; Xu et al., 2022; Yang et al., 2021, 2020). Therefore, we hypothesize that *bmp6* could be used to generate new cyprinid strains without IMBs by gene editing.

In this study, we generated a new crucian carp (*C. auratus*) strain without IMBs (hereafter WUCI strain) by knocking out *bmp6* and demonstrated that knockdown of *bmp6* did not affect the growth, and nutritional quality of the strain; we also conducted a metabolomics analysis of muscle to explore the effects of *bmp6* on substance synthesis and metabolism. This study provided an excellent germplasm for the cyprinids aquaculture industry and a useful molecular tool for eliminating IMBs in other cyprinids.

## 2. Materials and methods

### 2.1 Experimental fish

The diploid crucian carp (*C. auratus*, 2n=100) used in this study were cultured in the Hulan Experimental Station of HRFRI. Parental fish were reared in a 700m^2^ pond and injected with oxytocin drugs, including luteinizing hormone-releasing hormone (LRH-2, 4ug/kg body weight); domperidone (DOM, 1mg/kg body weight), and human chorionic gonadotropin (HCG, 100 U/kg body weight) for spawning. The fertilized eggs were incubated in 24cm×20cm×24cm incubation tanks at 22°C*∼*23°C. After hatching, larvae were fed 4 times/day of Artemia until they grew up to 2cm-3cm in body length; the larvae were moved to a 500m^2^ pond and fed artificial feed 2 times/day. The rear condition of F_0_, F_1_, F_2_, and F_3_ generation in the present study was the same as the above description.

### 2.2 *Bmp6* knockout

Because the crucian carp underwent the fourth round of genome duplication event(Chen et al., 2019), there are two orthologs of *bmp6* in the crucian carp genome. To identify the two orthologs, the cDNA and protein sequence of zebrafish *bmp6* were aligned to the crucian carp genome (Chen et al., 2019) using Blast tools, and the two orthologs were validated by the annotation of Ensembl(http://www.ensembl.org), which were ENSCARG00000018439(*bmp6a*) and ENSCARG00000056616(*bmp6b*).

The *bmp6a* and *bmp6b* knockout were carried out according to the method described by Xu et al. (2022) and described as follows. The target sites for CRISPR/Cas9 knockout were designed using the CHOPCHOP service (Labun et al., 2019) (Supplementary Table S1). The sgRNA was synthesized using the NEB HiScribe T7 Fast and Efficient RNA Synthesis Kit (NEB E2050S, MA, USA). After mixing 5μM~10μM sgRNAs of *bmp6a* and *bmp6b* and 5μM Cas9 protein (M0646, NEB, MA, USA), 1nL mixture was micro-injected into single-cell embryos. The embryos were incubated at 22~23°C as described above.

### 2.3 Mutants screening

After four months of culture, fish were labeled with PIT tags, and fin clips were collected for DNA extraction. The fin clips were lysed with 100μl solution consisting of 0.5 mg/ml proteinase K, 10 mmol/L pH 8.0 1 M Tris, 50 mmol/L KCl, 0.3% Tween 20, 0.3% NP 40 at 55°C for 6 h and 98°C for 10 min. After lysis, the lysate was mixed and centrifuged at 1000-2000 rpm for 2 min, and the supernatant was taken for PCR amplification.

The method for screening mutants is illustrated in Fig. 2(c). In this procedure, PCR amplification was performed using the method described by Tong et al. (2018) and was briefly described as follows. Two pairs of primers were used for PCR amplification, one of which was the target primer for amplifying the target fragment (Supplementary Table S2), and the other was the index primer for indexing samples. A 2-step amplification was performed: firstly, the PCR was carried out using 10 μ L volume, including genomic DNA supernatant 1 μ L, 1 μ mol/L forward and reverse target primers 0.5μL, 2× Dream Taq Master Mix (Thermo Fisher, CA, USA) 5μL, the PCR was performed with 95°C for 3 min, followed by 10cycles of 95°C 30 s, 60°C 30 s, 72°C 30 s; 72°C for 2 min; the second PCR amplification was performed by adding 0.5μL of 5 μmol/L forward and reverse index primers to the first step PCR product, and the PCR program was set to 95°C for 2 min; 6 cycles of 95°C 30 s, 58°C 30 s, 72°C 30 s; and 15 cycles of 95°C 30 s, 72°C 30s, finally 72°C 2min.

After PCR amplification, PCR products were mixed in equal amounts, DNA sequencing libraries were constructed using the TrueSeq library prepare kit, and 300bp Pair-End sequencing was performed using the Illumina MiSeq sequencing platform. The data obtained by MiSeq were first used to identify the loci and samples with the method described by Tong et al. (2018), followed by analysis with CRISPResso2 (Fiume et al., 2019) to screen the mutants and calculate the somatic mutation rate of each sample.

### 2.4 Intermuscular bone observation

We used skeleton staining, X-ray, and micro-CT to observe IMBs. Skeleton was stained using Alizarin red with the method described by Yang et al. (2020) and observed using the Zeiss Stereo Discovery V8 system (Carl Zeiss, Jena, Germany), the X-ray observation was performed using Faxitron MX-20 X-ray System(Faxitron, MA, USA), and the Quantum GX micro-CT Imaging System (PerkinElmer, MA, USA) with 70 KV, 88μA, and 14 min scan time was used for micro-CT. Around 50dph~60dph fish were used for bone staining, and four-month-old fish was used for X-ray and micro-CT scan. The counts of IMBs were checked using the images from bone staining and the X-ray.

### 2.5 Growth and reproductive performance evaluation

After one month of culture in a pond, 600 fingerlings of WUCI strain with BW of 4.6±0.42g and 600 fingerlings of wild-type crucian carp with BW of 4.2±0.55g were raised in a 500m^2^ pond. During the culture, the SL and BW were measured each month. At the age of 4 months, 50 individuals of WUCI strain and wild-type crucian carp were randomly chosen to measure the BW and SL. Furthermore, the gonad of 30 individuals from each group was dissected for weighting and histological examination according to Gan et al. (2021). The absolute fecundity and relative fecundity were calculated with the method described by Franěk et al.(2021).

### 2.6 Flesh quality evaluation

Thirty 4-month-old fish from each group were selected to evaluate flesh quality. For the texture analysis, the dorsal muscle with a size of 1cm^3^ and 2cm*1cm*1cm was dissected to measure the hardness, cohesiveness, springiness, gumminess, chewiness, and shear force using a TMS-Pro texture meter (Food Technology Corporation, VA, USA). The remained muscle tissues were dissected to measure the nutrient components, and muscles from 10 individuals were pooled; 3 pooled samples of each group were tested. Moreover, the content of trace elements was measured using dorsal muscle and liver tissues from 5 fish of each group. Additionally, dorsal muscles from 5 fish of each group were dissected for histological analysis according to Yang et al.(2021).

The nutrient components, including proximate compositions (moisture, crude ash, crude protein, and crude lipid), amino acid compositions, fatty acid compositions, and vitamin compositions (A, B1, B2, D, and E) were measured in Qingdao Sci-tech Innovation Quality Testing Co., Ltd. according to the national standards of China. The contents of trace elements, including Mg, Mn, Fe, Zn, Cu, Se, and K, were measured using inductively coupled plasma-mass spectrometry (ICP-MS, Agilent 7500cx, USA).

### 2.7 RT-qPCR of *bmp6a* and *bmp6b*

Ten organs, including intestine, liver, dorsal muscle, caudal muscle, brain, spleen, skin, gill, kidney, and heart from 6 wild-type crucian carp at 4-month-age were dissected for total RNA extraction using Trizol (Thermo Fisher, CA, USA). RNA quality was measured using Nanodrop 8000 (Thermo Fisher, CA, USA), and the quantity of RNA was measured using the Qubit3 kit (Thermo Fisher, CA, USA). The RNA was reverse-transcribed with the Revert Aid™ First-Strand cDNA Synthesis Kit (Thermo Fisher, CA, USA). The qPCR was analyzed with primers listed in Supplementary Table S3, and *β-actin* was used as the reference gene (Dharmaratnam et al., 2021). The RT-qPCR was carried out using 10 μL volume, including 1 μL of 50ng/μL cDNA, 0.5μL of 10μM forward and reverse primer, 5μL of 2× Luna Universal qPCR Master Mix (NEB, MA, USA). The amplification program was set as follows: 95 °C for 15s, followed by 40 cycles of 95 °C 15s and 60 °C for 30s. The qPCR was performed using the QuantiStudio 6 Flex Real-Time PCR System (Thermo Fisher, CA, USA).

### 2.8 RNA fluorescence in situ hybridization

We used the caudal muscle tissues of 60dph wild-type crucian carp to detect the spatial expression of *bmp6a* and *bmp6b*. The RNA fluorescence in situ hybridization was carried out with the SABER-FISH protocol according to Kishi et al.(2019, 2018). The probes were designed by the method proposed by Beliveau et al. (Beliveau et al., 2018) and listed in Supplementary Table S4. the mRNA of *bmp6a* and *bmp6b* was detected using Alexa Fluor 532 probes, and the nucleus was stained using DAPI dye (Thermo Fisher, CA, USA). The cryosections were observed using a fluorescence microscope (OLYMPUS BX53, Japan), and the images were processed by ImageJ.

### 2.9 Metabolomics analysis

The dorsal muscle tissues of 15 fish from each group were dissected and frozen in liquid nitrogen for metabolomics analysis. The tissues were grounded with liquid nitrogen, and after resuspending with prechilled 80% methanol by vortex, the samples were incubated on ice for 5min and were centrifuged for 20min at 15000g and 4°C; the supernatants were diluted to a final concentration of 53% methanol by LC-MS grade water and transferred to fresh Eppendorf tubes. After centrifugation at 15000g and 4°C for 20min, the supernatant was subjected to the LC-MS/MS system. The UHPLC-MS/MS analyses were carried out using a Vanquish UHPLC system (Thermo Fisher, Germany) coupled with an Orbitrap Q Exactive HF-X mass spectrometer (Thermo Fisher, Germany) in Novogene Co., Ltd. (Bejing, China).

The original file obtained by UHPLC-MS/MS was imported into Compound Discoverer 3.1 (CD3.1) software for s pectrogram processing with parameters of retention time tolerance of 0.2 min, an actual mass tolerance of 5ppm, signal intensity tolerance of 30%, and signal/noise ration of 3. After normalization to the total spectral intensity, the normalized peak intensities were used to predict the molecular formula based on additive ions, molecular ion peaks, and fragment ions. The databases of mzCloud(https://www.mzcloud.org), mzVault, and MassList database were used to match the peak and obtain accurate qualitative and relative quantitative results.

The quantitative matrix of metabolites was analyzed by multivariate statistical analysis, including principal component analysis (PCA) and partial least squares discriminant analysis (PLS-DA) to reveal the differences in metabolic patterns among different groups. The univariate analysis (Student’s t-test) was used to calculate the statistical significance, the metabolites with P-values<0.05, variable importance in the projection (VIP) value >1.0, and fold-change(FC) of WUCI vs. wild-type ≥1.5 or ≤0.667 were considered to differential metabolites. After annotating with KEGG, HMDB (https://hmdb.ca/metabolites), and LIPIDMaps database (http://www.lipidmaps.org), pathway analysis was carried out using MetaboAnalyst 5.0(Pang et al., 2022)

### 2.10 Statistical analysis

Student’s t-test was used to test the significance of the difference in body length, body weight, and muscle texture structure. And Wilcoxon Rank Sum Test was used to test the significance of the difference in reproductive performance, results of histological analysis, and content of elements and nutrients.

## 3. Results

### 3.1 Expression pattern of duplicated *bmp6* in crucian carp

*Bmp6* has been indicated to co-express in the myoseptum with osteoblast marker gene *sp7*, tendon progenitor marker gene *scxa*, and tenocyte marker gene *tnmd* and *xirp2a* (Xu et al., 2022). However, *bmp6* had two orthologs (*bmp6a*: ENSCARG00000018439; *bmp6b*: ENSCARG00000056616) in the crucian carp genome caused by the fourth whole-genome duplication event (Fig. 1(a))(Chen et al., 2019). To explore whether the two duplicated copies had a similar function to the development of IMBs in crucian carp, we investigated the expression levels in 9 tissues using RT-qPCR; *bmp6a* had a high expression level in dorsal muscle, caudal muscle, and skin; *bmp6b* had a high expression level in liver and gill (Fig. 1(a), (c)); Supplementary Figs. S1, S2). The spatial expression of *bmp6a* and *bmp6b* in caudal muscle detected by RNA fluorescence in situ hybridization (RNA-FISH) showed that both orthologs were expressed in the myoseptum(Fig. 1(d-e); Supplementary Fig. S3). These results indicated that both orthologs might function in IMB’s development despite the expressed levels.

**Figure 1.**
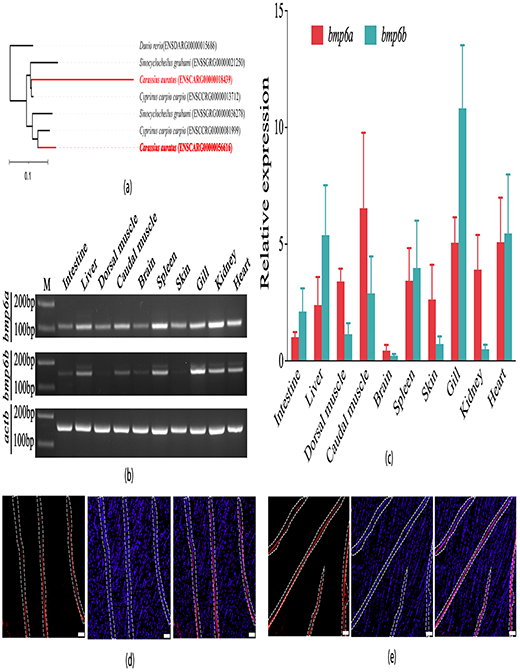
Expression of *bmp6a* and *bmp6b* in tissues of crucian carp. (a): the gene tree of *bmp6a* and *bmp6b*, the tree was extracted from Ensembl. (b): the expression of *bmp6a* and *bmp6b* in crucian carp tissues analyzed by qPCR, electrophoretic images were cropped from the original images in Supplementary Figs. S1, S2. M: molecular marker. (c): the relative expression levels of *bmp6a* and *bmp6b* in crucian carp tissues analyzed by real-time quantitative PCR. (d) and (e): spatial expression of *bmp6a* (d) and *bmp6b* (e) in caudal muscle, the white dashed lines indicate myospetum; mRNA of *bmp6a*(d) *and bmp6b*(e) were detected with Alexa Fluro 532 labeled probes (left), the nucleus was stained with DAPI dye (middle), and the merged images were shown in the right; scale bar: 100μm.

**Figure 2.**
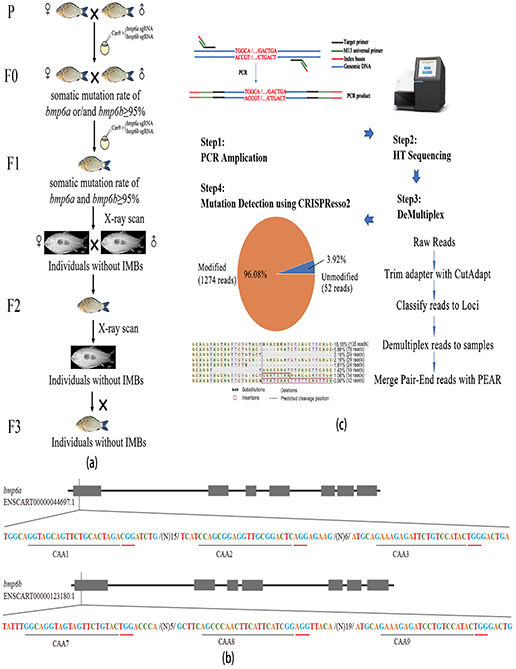
Pipeline for generating the new crucian carp strain without intermuscular bones. (a): the schematic diagram for constructing the new strain without intermuscular bones. (b): target sites of *bmp6a* and *bmp6b* for CRISPR/Cas9 knockout. (c): the schematic diagram for screening mutants.

### 3.2 Pipeline for *bmp6* gene editing in crucian carp

According to the expression in muscle, *bmp6a* and *bmp6b* should be knocked out simultaneously to obtain individuals without IMBs. In order to rapidly establish the WUCI strain, we designed a two-round gene knockout and mating strategy in this study (Fig. 2(a)). In this strategy, three target sites were designed in exon 1 of *bmp6a* and *bmp6b*, respectively (Fig. 2(b)). Firstly, sgRNAs of *bmp6a* and *bmp6b* were used for the first round of multiplex gene knockout to construct the F_0_ generation, and the mutants of F_0_ generation were screened by high-throughput sequencing using PCR products of target sites (Fig. 2(c)). The mutants with somatic mutation rates of more than 95% (*bmp6a* or *bmp6b*) were selected for natural mating, and the second round of multiplex gene knockout of *bmp6a* and *bmp6b* was performed in their fertilized eggs to construct the F_1_ generation. In the F_1_ generation, individuals with somatic mutation rates of more than 95% *bmp6a* and *bmp6b* were scanned by X-ray to screen individuals without IMBs, which were used to construct the F_2_ generation population by self-bred. Genotypes and counts of IMBs of the F_2_ generation were screened, and individuals without IMBs were used for reproducing the F_3_ generation by self-bred. Individuals of the F_3_ generation were used for evaluating the growth and fecundity performance, nutrient components, and metabolomics analysis.

### 3.3 Generation of a new strain without intermuscular bones

Using this pipeline, we carried out *bmp6* gene editing in crucian carp and generated the WUCI strain from 2019 to 2022(Fig. 3(a)). Among 668 F_0_ progenies, 511 mutants were screened, including 117 mutants with *bmp6a* and *bmp6b* mutated in somatic cells (Fig. 3(b) Supplementary Table S5). The somatic mutation rate of *bmp6a* and *bmp6b* in F_0_ mutants were 52.20%~96.10% and 20.80~99.80, respectively (Fig. 3(b), Supplementary Table S6), with the mean value of 72.1% and 75.74%, respectively. The percent of mutants with somatic mutation ratio *≥* 95% of *bmp6a* and *bmp6b* was 0.61% and 27.1%, respectively.

**Figure 3.**
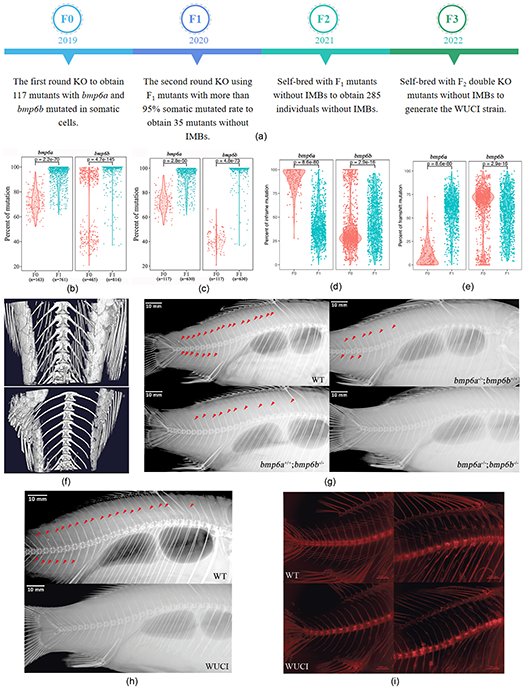
Observation of intermuscular bones in mutants of F_1_, F_2_, and F_3_ generations. (a): timeline of this study. (b): the somatic mutation distribution of mutants in the F0 and F_1_ generation. (c): the somatic mutation distribution of mutants with both genes mutated in the F_0_ and F_1_ generation. (d): in-frame mutation distribution of mutants in the F_0_ and F_1_ generation. (e): frame-shift mutation distribution of mutants in the F_0_ and F_1_ generation. (f): micro-CT images of a wild-type crucian carp (top) and a representative mutant without intermuscular bones in the F_1_ generation(bottom). (g): X-ray images of a wild-type crucian carp (top left) and representative mutants of single gene knockout homozygous line of *bmp6a*^−/−^ (top right), *bmp6b*^−/-^ mutant (bottom left), and *bmp6a*^−/-^/*bmp6b*^−/-^ (bottom right) in the F_2_ generation. (h): X-ray images of a wild-type crucian carp(top) and a representative mutant without intermuscular bones(bottom) in the F_3_ generation. (i): skeletal staining images of a wild-type crucian carp(top) and a representative mutant without intermuscular bones(bottom) in the F_3_ generation, 41 individuals were randomly chosen for bone staining, and no intermuscular bones were observed; top left, caudal skeleton; top right, dorsal skeleton. The scale bar in (g) and (h) was 10mm, and the scale bar in (i) was 2000μm.

After the knockout of *bmp6a* and *bmp6b* for the second time using F_0_ adults, 1130 F_1_ progenies were obtained, among which 947 mutants were screened, including 630 mutants with *bmp6a* and *bmp6b* mutated in somatic cells (Fig. 3(b), Supplementary Table S5). The somatic mutation rate of *bmp6a* and *bmp6b* in F_1_ mutants were 61.70%~100% and 36.70%~100%, respectively (Fig. 3(b), Supplementary Table S6), with the mean value of 93.4% and 97.25%, respectively. The percent of mutants with somatic mutation ratio *≥*95% of *bmp6a* and *bmp6b* was 64.65% and 89.58%, respectively.

A comparative analysis showed that the somatic mutation rates of *bmp6a* or *bmp6b* in the F_1_ mutants were significantly higher than those in the F_0_ mutants (*P*<0.0001, Fig. 3(b), Supplementary Table S5). The number of mutants with both genes mutated and the number of mutants with a somatic mutation rate of more than 95% in the F_1_ generation was also significantly higher than those in the F_0_ generation (*P*<0.0001, Fig. 3(c), Supplementary Table S6), and the somatic mutation rate of samples with both genes mutated in F_1_ mutants was also significantly higher than in F_0_ mutants (*P*<0.0001, Fig. 3(c), Supplementary Table S6). These results indicated that the two-round gene knockout strategy significantly increased the mutation rate and the number of samples with high mutation rates of both genes, resulting in the rapid establishment of homozygous mutated lines.

The in-frame and frameshift mutation rates of *bmp6a* and *bmp6b* in the F_0_ and F_1_ mutants were listed in Supplementary Table S6. The number of in-frame mutation of *bmp6a* in the F_0_ mutants were significantly higher than those in the F_1_ mutants, and the number of in-frame mutation of *bmp6b* in the F_1_ mutants were significantly higher than those in the F_0_ mutants (*P*<0.0001, Fig. 3(d)). The number of frame-shift mutation of *bmp6a* in the F_1_ mutants were significantly higher than those in the F_0_ mutants, and the number of frame-shift mutation of *bmp6b* in the F_0_ mutants were significantly higher than those in the F_1_ mutants (*P*<0.0001, Fig. 3(e)).

In the F_1_ generation, 270 progenies with 95% somatic mutation rate of *bmp6a* and *bmp6b* were screened, among which 35 mutants without IMBs were obtained using X-ray scanning (Fig. 3(f)), accounting for 12.96%; additionally, 110 mutants with more than 50% reduction of the count of IMBs were screened, accounting for 40.74%. The F_2_ population was constructed using the 35 F_1_ mutants without IMBs at one year old by self-bred; among 1500 F_2_ progenies, 285 individuals without IMBs were obtained, accounting for 19.00%, and 380 individuals with the count of IMBs reduced by more than 50%, accounting for 25.33% (Fig. 3(g)). Furthermore, sexually matured individuals without IMBs of the F_2_ generation at one year old were used to construct the F_3_ generation population by self-bred, and 41 individuals at 60dph (days post-hatched) were randomly chosen for bone staining, no IMB was found in staining (Fig. 3(h)). We also examined the IMBs by X-ray using 30 individuals at 4 months of age and approved the results (Fig. 3(i)). These results indicated that the WUCI strain was generated successfully.

The genotype analysis of F_1_, F_2_, and F_3_ generation showed that the WUCI strain was a multiple mutated alleles population. We randomly chose five individuals from 35 F_1_ mutants without IMBs to sequence the target site of *bmp6a* and *bmp6b*, and the results showed that 21 mutated alleles in somatic cells were observed in *bmp6a* (Supplementary Fig. S4-S5) and 11 mutated alleles in somatic cells were observed in *bmp6b* (Supplementary Fig. S6-S7), a frame-shift mutation occurred at the around 23^rd^ amino acid of *bmp6a* and *bmp6b*(Supplementary Fig. S5, S7). The genotypes of F_2_ individuals without IMBs (bottom right in Fig. 3(g)) used for constructing the F_3_ generation showed that 9 mutated alleles were observed in *bmp6a* and 5 mutated alleles were observed in *bmp6b*(Supplementary Figure S8), all the mutation caused frame-shift at the around 23^rd^ amino acid in *bmp6a* and the around 22^nd^ amino acid of *bmp6b*. Additionally, we carried out the knockout of a single gene (*bmp6a* or *bmp6b*) and checked the genotypes (Supplementary Figure S9) and phenotypes of the homozygous lines (*bmp6a*^−/−^;*bmp6b*^+/+^ at the top right of Fig. 3(g) and *bmp6a*^+/+^;*bmp6b*^−/−^ at the bottom left of Fig. 3(g)), the results showed that the knockout of *bmp6a* or *bmp6b* caused the partial deletion of IMBs in homozygous mutants (top right and bottom left in Fig. 3(g)). We also randomly genotyped 5 individuals of the F_3_ generation and observed 5 mutated alleles in *bmp6a* and 4 mutated alleles in *bmp6b*, which were inherited from the F_2_ parents (Supplementary Fig. S10).

### 3.4 Growth of the new strain

For evaluating the growth performance of the WUCI strain, 600 one-month-old fingerlings of WUCI strain with body weight (BW) around 4.6±0.42g and body length(SL) around 4.94±0.38mm and 600 one-month-old fingerlings of wild-type crucian carp with BW around 4.2±0.55g and SL approximately 4.79±0.35mm were cultured for three months at a 500m^2^ pond, the BW and SL were measured each month. The relationship between BW and SL of the WUCI strain was similar to that of the wild-type crucian carp (Supplementary Fig. S11). The comparisons showed that the SL and increment of SL per month of the WUCI strain were significantly higher than that of wild-type crucian carp at 2, 3, and 4 months of age(Fig. 4(a-b), Supplementary Table S7); there was no significant difference in specific growth rate(SGR) of SL per month between the WUCI strain and wild-type crucian carp at 2,3months of age, whereas, a significant difference(*P*<0.001) in SGR of SL at four months of age was observed(Fig. 4(c), Supplementary Table S7). The growth of BW was similar to that of SL (Fig. 4(d-f), Supplementary Table S7). Based on the above analysis results, it can be concluded that the WUCI strain at 2~4-month-old grew faster than the wild-type crucian carp.

**Figure 4.**
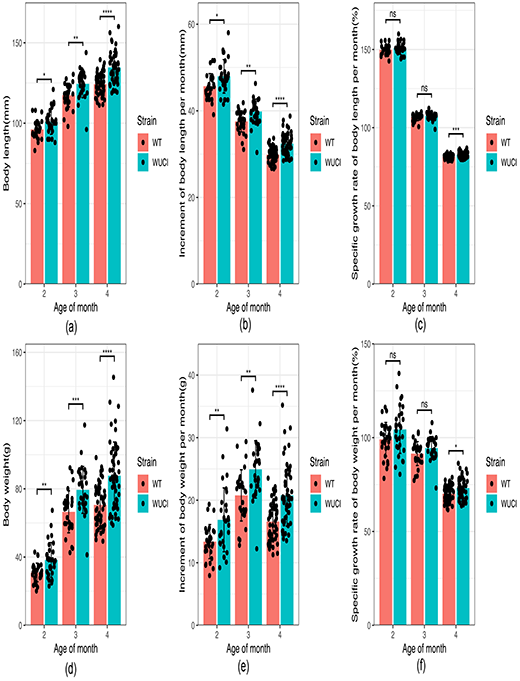
Growth performance of the new strain without intermuscular bones and the wild-type crucian carp. The body length (SL, a) and body weight (BW, b) were measured at the age of 2, 3, and 4 months, the increment of SL(d) and BW(e) per month and the specific growth rate per month of SL(c) and BW(f) was calculated. The summary of BW and SL were listed in Supplementary Table S7. The significant difference was tested using Student’s t-test. WT: the wild-type crucian carp; WUCI: the new strain without intermuscular bones; *: *P*<0.05; **: *P*<0.01; ***: *P*<0.001; ****: *P*<0.0001; ns: no significance.

### 3.5 Nutrient components of the new strain

The nutrient components in muscle tissue of 4-month-old WUCI strain and wild-type crucian carp were determined. We analyzed the proximate composition, total saccharide, vitamins, fatty acids, and amino acids (Supplementary Table S8-11). Seventeen kinds of amino acids and 17 types of fatty acids were determined. Among the five vitamins (A, B1, B2, D, and E) we measured, vitamins A and D were not detected. The comparison results showed that there was no significant difference in the contents of nutrient components mentioned above, as well as essential amino acid (EAA), umami amino acid (UAA), essential fatty acid (EFA), polyunsaturated fatty acid (PUFA), Omega3, and Omega6 (Fig. 5, Supplementary Table S8-11). We also measured seven elements in the muscle and liver tissues, including Mg, Mn, Fe, Cu, Zn, Se, and K. The significance test showed that the contents of Fe and Se in the liver of the WUCI strain were significantly higher than those of the wild-type crucian carp. In contrast, the contents of all elements in the muscle did not show a significant difference (Fig. 6, Supplementary Table S12).

**Figure 5.**
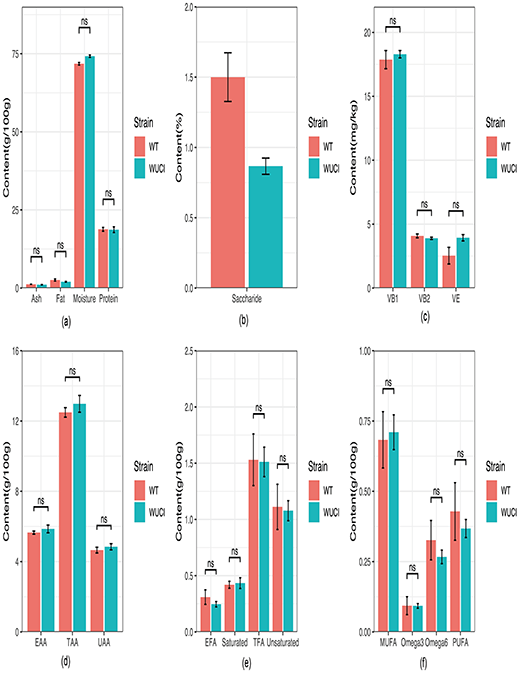
Nutrient quality of the new strain without intermuscular bones and the wild-type crucian carp. Proximate composition(a), total saccharide(b), vitamins(c), amino acid composition(d), and fatty acid (e, f) were determined. The data used to draw these figures were shown in Supplementary Table S8-S11. Wilcoxon test was used to test the significant levels (n=3). WT: the wild-type crucian carp; WUCI: the new strain without intermuscular bones; Ash: crude ash; Fat: crude fat; Protein: crude protein; Saccharide: total saccharide; VB1: vitamin B1; VB2: vitamin B2; VE: vitamin E; EAA: essential amino acid; TAA: total amino acid; UAA: umami amino acid; EFA: essential fatty acid; Saturated: saturated fatty acid; TFA: total fatty acid; Unsaturated: unsaturated fatty acid; MUFA: monounsaturated fatty acid; PUFA: polyunsaturated fatty acid; Omega3: omega3 fatty acid; Omega6: omega6 fatty acid; ns: no significance.

**Figure 6.**
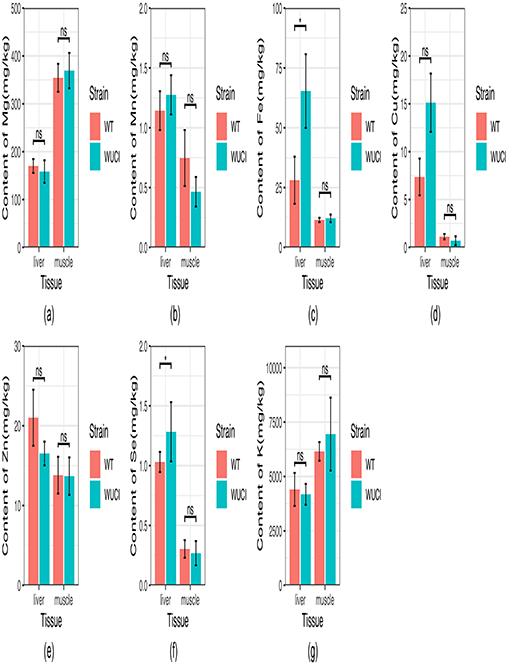
Content of 7 elements in the new strain without intermuscular bones and the wild-type crucian carp. The contents of Mg(a), Mn(b), Fe(c), Cu(d), Zn(e), Se(f), and K(g) in the new strain, and the wild-type crucian carp were determined using the dorsal muscle and liver tissues. The data used to draw these figures were listed in Supplementary Table S12. Wilcoxon test was used to test the significant levels(n=5). WT: the wild-type crucian carp; WUCI: the new strain without intermuscular bones; *: *P*<0.05; ns: no significance.

### 3.6 Muscle texture and structure of the new strain

Muscle texture and structure are important to fish flesh quality. The muscle texture analysis showed that the shear force of the WUCI strain was significantly higher than those of wild-type crucian carp(*P*<0.01) (Fig. 7(a), Supplementary Table S13). In contrast, the hardness of the WUCI strain was significantly weaker than those of wild-type crucian carp(*P*<0.05) (Fig. 7(b), Supplementary Table S13). Furthermore, the two groups had no significant difference in cohesiveness, springiness, gumminess, and chewiness (Fig. 7(c-f), Supplementary Table S13). The PCA analysis result of muscle texture showed that most samples of the WUCI strain were clustered together with those of the wild-type strain and not well distinguished (Supplementary Fig. S12), indicating that no significant difference occurred between the two groups. We also carried out a histological analysis of muscle tissues, and the results showed that the muscle fiber area and density had no significant differences (*P*>0.05) between the WUCI strain and the wild-type crucian carp (Fig. 7(g-i), Supplementary Table S13). These results indicated that the knockout of *bmp6a* and *bmp6b* and the lack of IMBs did not significantly change the muscle texture and structure of the WUCI strain.

**Figure 7.**
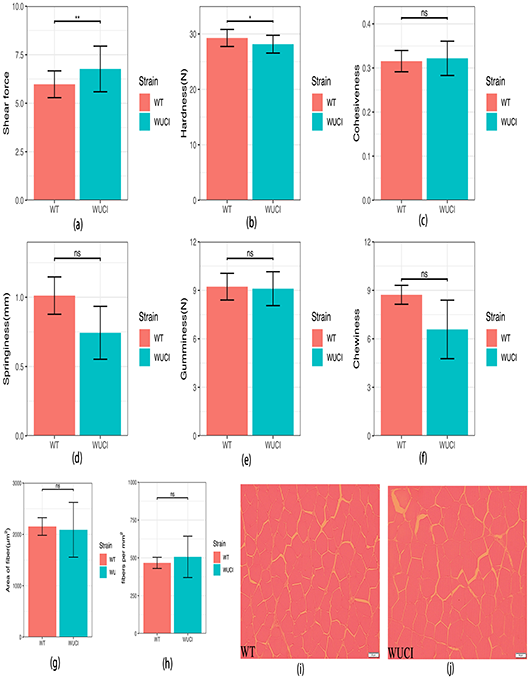
Muscle texture and structure of the new strain without intermuscular bones and the wild-type crucian carp. The shear force(a), hardness(b), cohesiveness(c), springiness(d), gumminess(e), and chewiness(f) were tested using a texture meter; Student’s t-test was used to test the significant levels(n=30). The histological sections were carried out using dorsal muscle in the wild-type crucian carp(i) and the new strain(j), and the area(g) and density(h) of muscle fibers were calculated using the histological images with the ImageJ program, Wilcoxon test(n=5) was used to test the significant levels. The data used to draw the figures were shown in Supplementary Table S13. The scale bar in (i) and (j) was 50μm. WT: the wild-type crucian carp; WUCI: the new strain without intermuscular bones; *: *P*<0.05; **: *P*<0.01; ns: no significance.

### 3.7 Reproductive performance of the new strain

The comparison analyses showed no significant difference in the weight of ovary and ovarian GSI (Gonadosomatic index, gonad weight/body weight ×100) between the 4-month-old WUCI strain and wild-type crucian carp(*P*>0.05) (Fig. 8(a-b); Supplementary Table S14). The weight of the testis and GSI of the WUCI strain were significantly higher than those of wild-type crucian carp (*P*<0.01 and *P*<0.05, respectively) (Fig. 8(a-b); Supplementary Table S14). Moreover, no significant difference between the two groups was observed in the absolute female fecundity (total quantity of produced eggs by females) and relative fecundity (the number of eggs per gram of body weight) (*P*>0.05) (Fig. 8(c-d); Supplementary Table S14). The gonad histological analysis showed that the development of the ovary and testis of the WUCI strain was similar to that of the wild-type crucian carp (Fig. 8(e-f)). These results suggested that the *bmp6a* and *bmp6b* knockout and the absence of IMBs did not significantly affect the gonad development and the fecundity of the WUCI strain.

**Figure 8.**
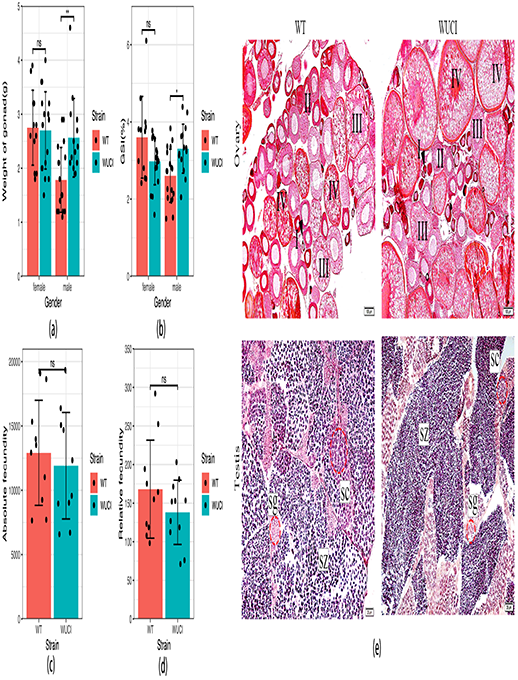
Reproductive performance of the new strain and the wild-type crucian carp. The weight of gonad(a), gonad index (GSI, b), female absolute fecundity(c), and relative fecundity(d) were compared between the 4-months-old individuals of the new strain and the wild-type crucian carp. The data used to draw the figures were shown in Supplementary Table S14; the Wilcoxon test was used to test the significant levels. (e), histological sections of gonads in the wild-type crucian carp and the new strain, the scale bar in ovary and testis tissues was 100μm and 20μm, respectively. WT: the wild-type crucian carp; WUCI: the new strain without intermuscular bones; I(black arrowhead), II, III, and Ⅳ represent the developmental phases of oocytes; sg: Spermatogonium; sc: spermatocytes; SZ: Spermatozoa; *: *P*<0.05; ns: no significance.

### 3.8 Metabolome analysis

The peaks extracted from all samples and QC were analyzed by PCA, and the results showed that the QC samples in the positive and negative ion modes were clustered together (Fig. 9(a-b)), indicating that the experiment was robust. The PLS-DA model of two groups was established, and 7-fold cross-validation was performed to characterize the model’s stability and reliability (Fig. 9(c-d)). A total of 863 metabolic ion peaks were extracted, including 546 positive ion peaks and 317 negative ion peaks. Using the parameters of VIP > 1.0, FC > 1.5 or FC < 0.667, and P-value< 0.05, 107 metabolites in the positive ion mode had significant differences, including 28 up-regulated and 79 down-regulated metabolites in the WUCI strain (Supplementary Table S15); in the negative ion mode, 124 metabolites had significant differences, including 5 up-regulated and 119 down-regulated metabolites in the WUCI strain (Supplementary Table S16). These up-regulated metabolites included some metabolites that are beneficial to the organism, such as β-Nicotinamide mononucleotide, DL-Glutamine, Hypoxanthine, Traumatic acid, Thiamine pyrophosphate, and so on (Fig. 9(e-l)). KEGG pathway enrichment analysis showed that 46 metabolite pathways were enriched, including Thiamine metabolism, Glutathione Metabolism, Nicotinate and Nicotinamide Metabolism, amino acids metabolism pathways, etc. (Fig. 9(m), Supplementary Table S17, Supplementary Fig. S13).

**Figure 9.**
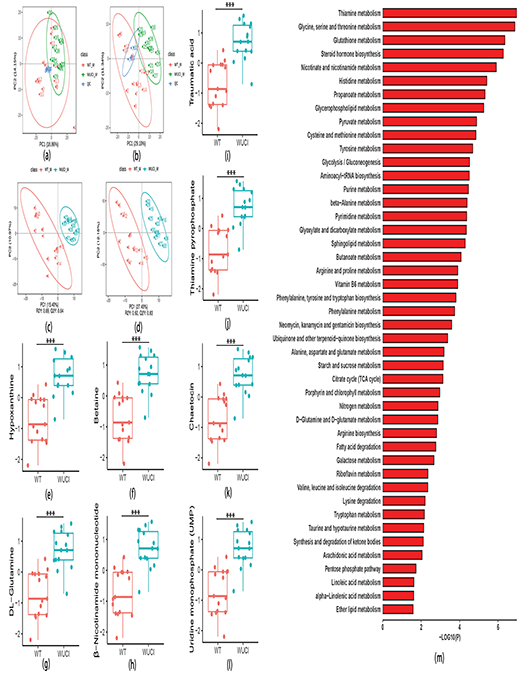
Metabolomics analysis of the new strain and the wild-type crucian carp. The PCA analysis of the metabolites in positive ion mode(a) and negative ion mode(b); The PLS-DA analysis of the metabolites in positive ion mode(c) and negative ion mode(d); (e-l): up-regulated metabolites in the WUCI strain. (m): the enriched pathways of metabolites in muscle tissue of the new strain analyzed by MetaboAnalyst 5.0. WT_M: muscle of wild-type crucian; WUCI_M: muscle of the new strain without intermuscular bones. \*\*\**:P*<0.001

## 4. Discussion

The cyprinids aquaculture industry requires varieties with high flesh quality, and the reduction of IMBs is one of the main targets for improving the flesh quality (Gjedrem, 1997; Gui et al., 2021; Sun and Zhu, 2019). Although researchers utilized many approaches to deplete cyprinids IMBs, little progress was obtained until the eruption of the gene editing technology. In our previous study(Xu et al., 2022), we characterized *bmp6* as an essential gene to the development of IMBs in zebrafish and obtained a zebrafish strain with IMBs completely absent by knockout *bmp6*, in this study, we generated a new strain without IMBs by knocking out the two orthologs, *bmp6a* and *bmp6b*. This strain has expected performances of growth, reproduction, nutrient value, and muscle texture and structure; significantly, its muscle enriched some beneficial metabolites for human health. According to our knowledge, it is the first report that a farmed cyprinid strain without IMBs, which could be stably inherited, was obtained in the world.

There are two orthologs of *bmp6* in crucian carp; according to the expression in the muscles, it is supposed that the two orthologs should be knocked out simultaneously to obtain mutants with complete depletion of IMBs. To test this hypothesis, we generated two additional homozygous lines by knocking out a single gene, *bmp6a* or *bmp6b*, the phenotypes of homozygous mutants of *bmp6a*^−/−^ or *bmp6b*^−/−^ showed that the knockout of *bmp6a* or *bmp6b* caused the partial deletion of IMBs in mutants, whereas the double knockout of *bmp6a* and *bmp6b* caused the complete elimination of IMBs in the F_2_ generation which also approved by the phenotypes of the F_3_ generation because the F_3_ generation was established by self-bred with *bmp6a*^−/-^;*bmp6b*^−/-^ F_2_ adults.

In general, multiplex gene knockout homozygous strains were obtained by sequential knocking out a single gene or crossing with homozygous lines with a single gene knockout, which is difficult and time-consuming (around 3~4 generations to get a homozygous line). We established a two-round knockout strategy to obtain homozygous mutants with the simultaneous knockout of *bmp6a* and *bmp6b*. This strategy contains three aspects: firstly, we used three guides to knock out *bmp6a* or *bmp6b*. Wu et al. (2018) proved that the simultaneous use of multiple guides could significantly increase the efficiency of functional disruption; we adopted this strategy and obtained mutants in more than 70% of the F_0_ generation (Supplementary Table S5, S6). Secondly, the mutants with more than 95% mutation rate of somatic cells were chosen to reproduce the next generation. Generally, the mutation in germ cells was unknown, and we supposed that the gene in germ cells should be mutated if the somatic mutation rate was more than 95%, if we used the fertilized eggs from those F_0_ adults to generate the F_1_ generation by the second knockout, it could significantly increase the simultaneous mutated probabilities of *bmp6a* and *bmp6b* because the mutation of the F_1_ progenies was the accumulation of the mutation inherited from the F_0_ germ cells and the mutation caused by the second knockout. This assumption was evidenced by our results, in which the somatic mutation ratio in the F_1_ generation was significantly higher than that in the F_0_ generation (Fig. 3(b-c); Supplementary Table S5, S6). Thirdly, the mutants were screened using high-throughput sequencing technology. Genome editing in cultured fishes needs to screen many individuals to obtain as many mutants as possible, which is labor- and time-consumption by traditional methods, such as gel electrophoresis, Sanger sequencing, high-resolution melt curves, etc. Based on our previous study (Tong et al., 2018), we developed a mutant screening method using high-throughput sequencing of PCR products by amplifying the target fragments using two pairs of primers, one of which was used to amplify the fragment containing the gene editing target sites and the other pair was used to append the index bases of the sample to the PCR product, after sequencing, the data were demultiplexed to individuals and knockout targets which were used to screen the mutants by CRISPResso2 (Fiume et al., 2019). Using this method, we can obtain the mutants in a time- and cost-efficient way. This strategy has an advantage, i.e., it could reduce the time to establish a new strain with both genes mutated from 3~4 generations to 2 generations; however, the disadvantage is that the strain has multiple alleles (Supplementary Figs. S4-S10).

Using the strategy described above, we obtained the mutants without IMBs in the F_1_ population and established the WUCI strain in two generations; the observation of IMB in the F_3_ progenies suggested that the trait was stable and inheritable; these results demonstrated that *bmp6* could be used to generate new cyprinids strains without IMBs by gene editing.

*Bmp6* is a member of the BMP family, and it regulates many biological processes, such as skeletal development(Wan and Cao, 2005), iron metabolism(Jr et al., 2009), female fertility (Campbell et al., 2009; Sugiura et al., 2010), and so on. Although the knockout of *bmp6* had no significant effects on growth and reproductive performance or the development of muscle and skeletal in zebrafish (Jian et al., 2019; Xu et al., 2022; Yang et al., 2021, 2020), it is unknown whether the knockout of *bmp6* affects the performance of the WUCI strain. Here, we comprehensively evaluated growth, reproduction, nutrient quality, muscle structure, and texture between the WUCI strain and the wild-type crucian carp. The results showed that the knockout of *bmp6a* and *bmp6b* did not affect the growth and reproductive performance of the WUCI strain, which was consistent with our previous studies in zebrafish (Jian et al., 2019; Xu et al., 2022; Yang et al., 2021, 2020). Growth is associated with the development of muscle and skeleton; the skeletal phenotype and the histological section of muscle showed that the musculoskeletal development was normal in the WUCI strain, implying that the function of *bmp6* in the musculoskeletal system might be compensated by others of the BMP family (Solloway et al., 1998). *Bmp6* is also involved in proliferation, steroidogenesis, and oocyte development in humans and mammals(Liu et al., 2019; Lochab and Extavour, 2017; Song et al., 2022; Sugiura et al., 2010; Zhang et al., 2019) functions in a specifies-specific way(Regan et al., 2018). Although the mRNA of *bmp6* was detected in oocytes and granulosa cells(Li and Ge, 2011) in zebrafish, there was no enough evidence to show the effect of *bmp6* on female fecundity. The present study and our previous studies showed that the knockout of *bmp6* did not significantly affect the reproductive performance in crucian carp and zebrafish, which implied that *bmp6* might not be the essential gene in the gonadal development of cyprinids, or the function of *bmp6* is compensated by other members of the BMP/TGE-β family; however, more studies are required. The detection of 7 elements in the liver and muscle showed that the content of Fe was higher in the WUCI strain than that in the wild-type crucian carp, which indicated that the iron overload in the liver of the WUCI strain caused by the dysfunction of *bmp6* (Jr et al., 2009), however, the content of Fe in the muscle of the WUCI strain did not show the significance compared to that of the wild-type crucian carp, which meant the deposited and transferred mechanism of Fe in muscle might be different in the liver.

In addition to evaluating nutrient quality by the traditional method, we performed further analysis of the metabolome. Metabolomics analysis results showed some pathways and metabolites were enriched in the muscle tissue of the WUCI strain, such as Nicotinate and nicotinamide metabolism pathway, Thiamine metabolism pathway, β-Nicotinamide mononucleotide, DL-Glutamine, Hypoxanthine, and Betaine. These pathways and metabolites are related to beneficial properties, including anti-aging, anti-oxidative, and anti-radiation damage effects (Fig. 9). Nicotinate and nicotinamide are precursors of the coenzymes nicotinamide adenine dinucleotide (NAD+) and nicotinamide adenine dinucleotide phosphate (NADP+) that are essential for the generation of coenzymes (Information, 2022). β-Nicotinamide mononucleotide is an important metabolite of the Nicotinate and nicotinamide metabolism pathway (Supplementary Fig. S13A). β-Nicotinamide mononucleotide is also a precursor of NAD+ and is considered a candidate compound for suppressing aging because it retained in the body for longer than Nicotinamide (KAWAMURA et al., 2016; Nadeeshani et al., 2021). Thiamin is involved in several cell functions, including energy metabolism and the degradation of sugars and carbon skeletons (Manzetti et al., 2014). Thiamine pyrophosphate is the active form of thiamine and a member of the Thiamine metabolism pathway (Supplementary Fig. S13B) (Zeece, 2020) deficiencies can lead to various forms of beriberi (Nohr et al., 2011). DL-Glutamine is the most abundant amino acid in the body. A recent study has shown that adding enteral L-glutamine to normal nutrition during the early stage of COVID-19 infection can shorten the hospital stay for patients (Cengiz et al., 2020). Hypoxanthine is an intermediate metabolite in the purine degradation system and a substrate for ATP synthesis which can reduce the severity of radiation damage by increasing the survival of endothelial cells from DNA damage and apoptosis in mice (Fujiwara et al., 2022). Betaine is lipotropic, an important human nutrient, and dietary betaine was utilized as an osmolyte and source of methyl groups, thereby helping maintain liver, heart, and kidney health(Craig, 2004).

## 5. Conclusion

*Bmp6* is a key gene that regulates the development of intermuscular bones. For the two orthologs of *bmp6* in crucian carp genome, we established a two-round gene knockout and mating strategy to rapidly generate the new crucian carp strain without IMBs by knocking out *bmp6*. We cultured the WUCI strain to the F3 generation and characterized the strain outperformed the wild-type crucian carp in growth at 4-month age and did not show significant differences in the reproductive performance and flesh quality. The metabolomics analysis of the muscle tissues showed that the WUCI strain significantly enriched some metabolites plays beneficial effects in anti-aging, anti-oxidant, and anti-radiation damage. This study is the first report that a farmed cyprinid strain without IMBs, which could be stably inherited, was obtained in the world. This strain provides excellent germplasm for the cyprinids aquaculture industry. Our gene editing strategy is a useful molecular tool for eliminating IMBs in other cyprinids and knocking out multiple orthologs rapidly.

## Acknowledgments

We greatly thank Professor Ying Li at the Second Affiliated Hospital of Harbin Medical University for scanning mutants with X-ray and thank Dr. Yi Zhou at Harvard Medical School for providing technical support in CRISPR/Cas9. We also thank Professor Yumei Chang, Chitao Li, and Gefeng Xu at HRFRI for their support in the histological section and muscle texture analysis.

## Ethics statement

In this study, all animal procedures were conducted according to the guidelines for the care and use of laboratory animals of Heilongjiang River Fisheries Research Institute (HRFRI), Chinese Academy of Fishery Sciences (CAFS). The studies in animals were reviewed and approved by the Committee for the Welfare and Ethics of Laboratory Animals of HRFRI, CAFS. The gene editing studies were approved by the Committee for Genetic Modification Organism (GMO) Safety Management (Approved ID: HRFRI-GMO-201801).

## CRediT authorship contribution statement

**Youyi Kuang:** Conceptualization, Methodology, Writing-Original Draft, Writing-Review & Editing, Supervision, Funding acquisition. **Xianhu Zheng**: Investigation, Resources. **Dingchen Cao**: Investigation, Resources. **Zhipeng Sun**: Investigation, Resources. **GuangxiangTong**: Investigation, Resources. **Huan Xu**: Writing-Original Draft, Writing-Review & Editing, Formal analysis. **Ting Yan**: Investigation, Funding acquisition. **Shizhan Tang**: Investigation, Formal analysis. **Zhongxiang Chen:** Investigation, Formal analysis. **Tingting Zhang**: Investigation,Visualization. **Tan Zhang:** Investigation,Visualization. **Le Dong**: Investigation,Visualization. **Xiaoxing Yang**: Investigation,Visualization. **Huijie Zhou**: Investigation,Visualization. **Weilun Guo**: Investigation,Visualization. **Xiaowen Sun**: Conceptualization.

