## Supplementary Figures for "Generate a new crucian carp (*Carassius auratus*) strain without intermuscular bones by knocking out *bmp6*"

**Supplementary figure legends**

**Supplementary Figure S1. Expression of *bmp6a*(top) and *bmp6b*(bottom) in crucian carp tissues.**

**Supplementary Figure S2. Expression of *β-actin* in crucian carp tissues.**

**Supplementary Figure S3. Spatial expression of *bmp6a*(A) and *bmp6b*(B) in caudal muscle.** White dashed lines and white arrowheads indicate myoseptum.

**Supplementary Figure S4. DNA Sequence alignments of *bmp6a* exon1 in $F_1$ mutants without intermuscular bones.**

**Supplementary Figure S5. Alignments of *bmp6a* protein sequences inferred with DNA sequences in $F_1$ mutants without intermuscular bones.**

**Supplementary Figure S6. DNA Sequence alignments of *bmp6b* exon1 in $F_1$ mutants without intermuscular bones.**

**Supplementary Figure S7. Alignments of *bmp6b* protein sequences inferred with DNA sequences in $F_1$ mutants without intermuscular bones.** WT represents wild-type, and the others represent mutants.

**Supplementary Figure S8. DNA sequences and protein sequences alignments of bmp6a exon1 and *bmp6b* exon1 in *bmp6a*$^{-/-}$;*bmp6b*$^{-/-}$ individuals without intermuscular bones in the $F_2$ generation.**

**Supplementary Figure S9. DNA sequences and protein sequences alignments of *bmp6a* exon1 in *bmp6a*$^{-/-}$;*bmp6b*$^{+/+}$ individuals(A-D) and *bmp6b* exon1 in *bmp6a*$^{+/+}$;*bmp6b*$^{-/-}$ individuals(E-H).**

**Supplementary Figure S10. DNA sequences and protein sequences alignments of exon1 of *bmp6a* and *bmp6b* in *bmp6a*$^{-/-}$;*bmp6b*$^{-/-}$ individuals of the $F_3$ generation.**

**Supplementary Figure S11. The relationship between body weight and body length.** A: the new strain without intermuscular bones; B: the wild-type crucian carp.

**Supplementary Figure S12. PCA analysis of muscle texture.** WT: wild-type crucian carp (WT), WUCI: the new strain without intermuscular bones.

**Supplementary Figure S13. Boxplot of metabolites in Nicotinate and Nicotinamide Metabolism(A) and Thiamine metabolism(B).**

**Supplementary table legends**

**Supplementary Table S1. Target sites of bmp6a and bmp6b for CRISPR/Cas9 knockout**

**Supplementary Table S2. Primer sequences for screening mutants**

**Supplementary Table S3. Primer sequences for RT-qPCR**

**Supplementary Table S4. Probe sequence for RNA-FISH of bmp6a and bmp6b**

**Supplementary Table S5. Mutants screening of $F_0$ and $F_1$ generation**

**Supplementary Table S6. Summary of somatic mutation ratio in $F_0$ and $F_1$ generation**

**Supplementary Table S7. Growth of the new strain**
**Supplementary Table S8. Proximate composition of the new strain and wild-type crucian carp**
**Supplementary Table S9. Vitamin composition of the new strain and wild-type crucian carp**
**Supplementary Table S10. Amino acid composition of the new strain and wild-type crucian carp**
**Supplementary Table S11. Fatty acid composition of the new strain and wild-type crucian carp**
**Supplementary Table S12. Element trace content of the new strain and wild-type crucian carp**
**Supplementary Table S13. Muscle texture of the new strain**
**Supplementary Table S14. Reproductive performance of the new strain**
**Supplementary Table S15. Metabolists with significant difference in the positive ion mode**
**Supplementary Table S16. Metabolists with significant difference in the negative ion mode**
**Supplementary Table S17. Enriched pathways of metabolites in muscle analyzed by MetaboAnalyst 5.0**

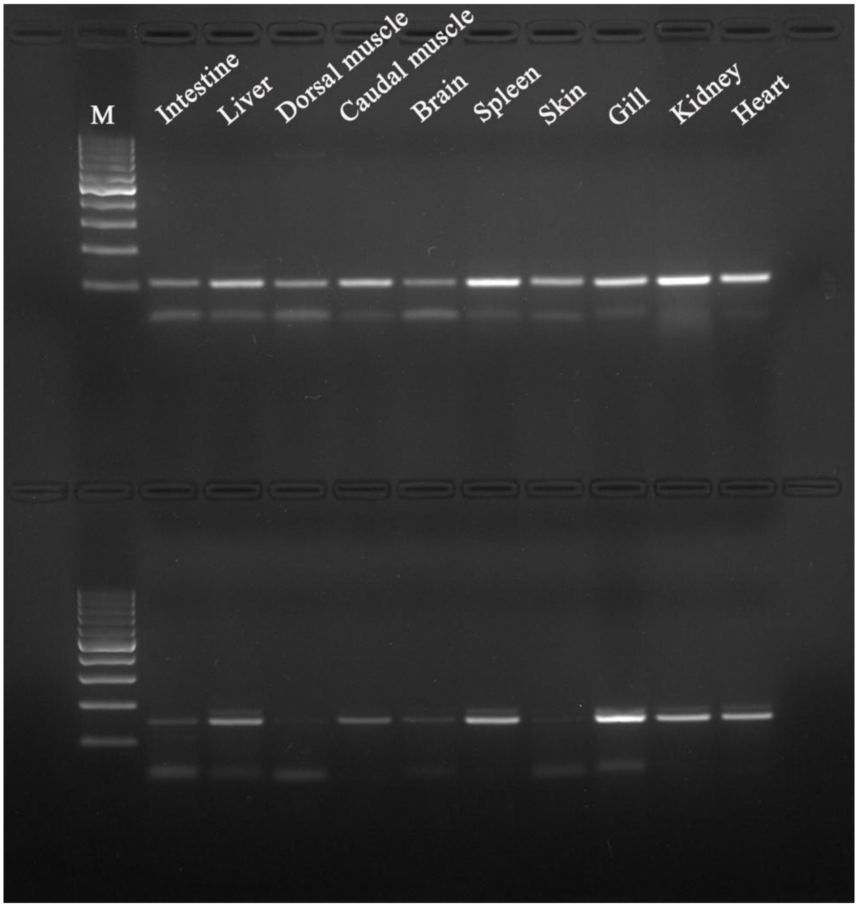


**Supplementary Figure S1. Expression of *bmp6a*(top) and *bmp6b*(bottom) in crucian carp tissues.** M: 100bp DNA marker.

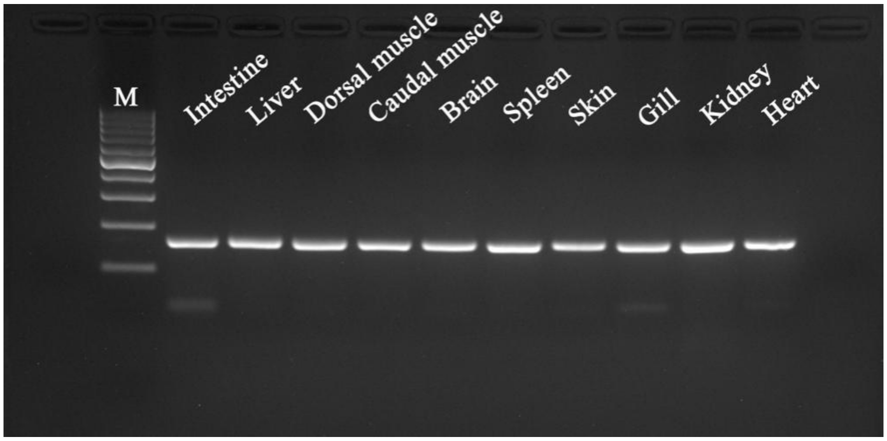


**Supplementary Figure S2. Expression of *β-actin* in crucian carp tissues.** M: 100bp DNA marker.

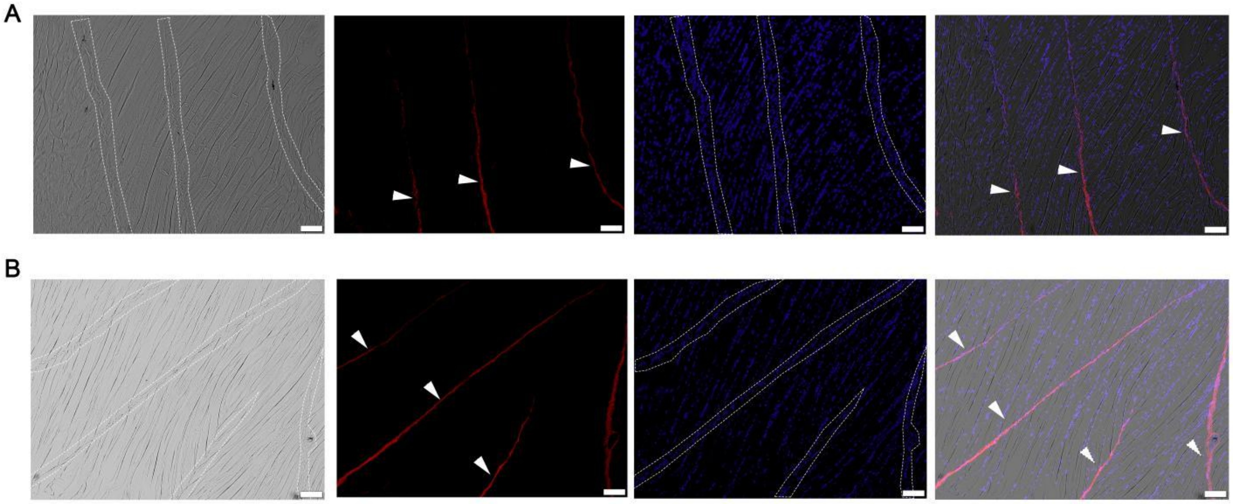


**Supplementary Figure S3. Spatial expression of *bmp6a*(A) and *bmp6b*(B) in caudal muscle.** White dashed lines and white arrowheads indicate myoseptum; scale bar: 100μm.

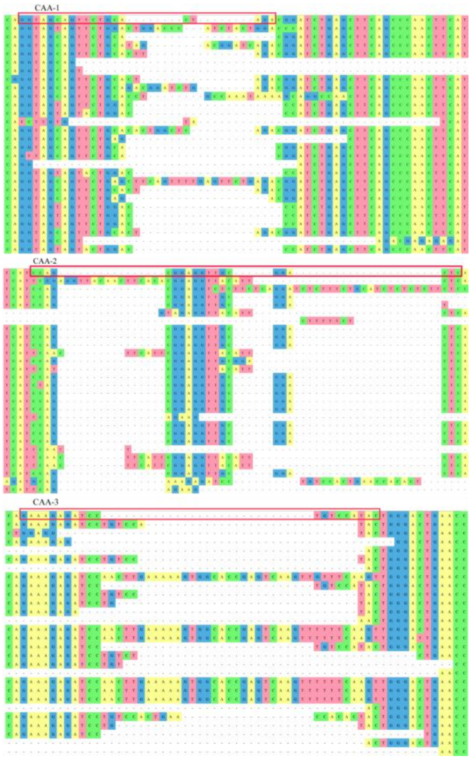

**Supplementary Figure S4. DNA Sequence alignments of *bmp6a* exon1 in $F_1$ mutants without intermuscular bones.** CAA-1, CAA-2, and CAA-3 were the target sites (marked with red box) for *bmp6a* knockout. WT represents wild-type, and the others represent mutants.

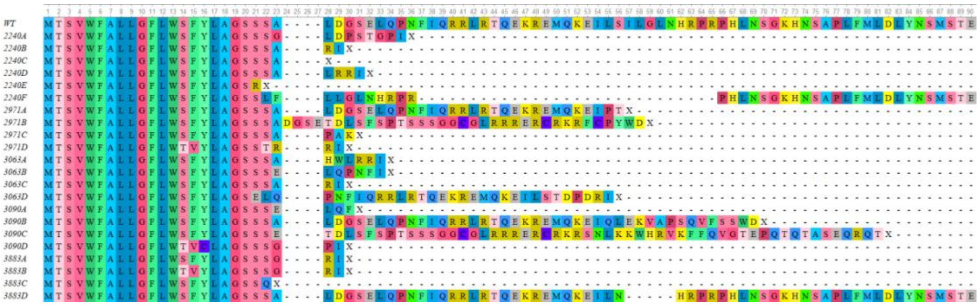

**Supplementary Figure S5. Alignments of *bmp6a* protein sequences inferred with DNA sequences in $F_1$ mutants without intermuscular bones.** WT represents wild-type, and the others represent mutants.

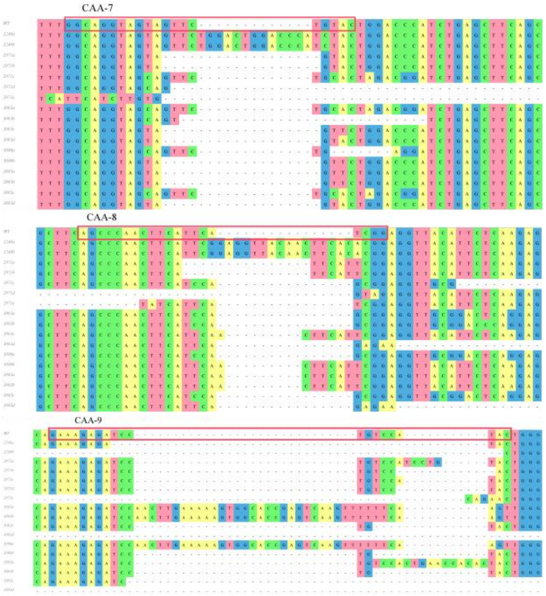


**Supplementary Figure S6. DNA Sequence alignments of *bmp6b* exon1 in F$_1$ mutants without intermuscular bones.** CAA-7, CAA-8, and CAA-9 were the target sites (marked with red box) for *bmp6b* knockout. WT represents wild-type, and the others represent mutants.

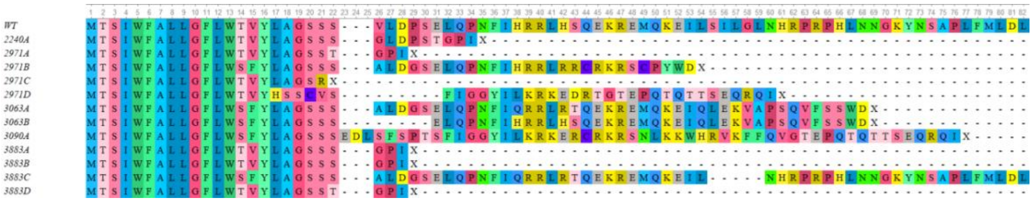


**Supplementary S7. Alignments of *bmp6b* protein sequences inferred with DNA sequences in F$_1$ mutants without intermuscular bones.** WT represents wild-type, and the others represent mutants.

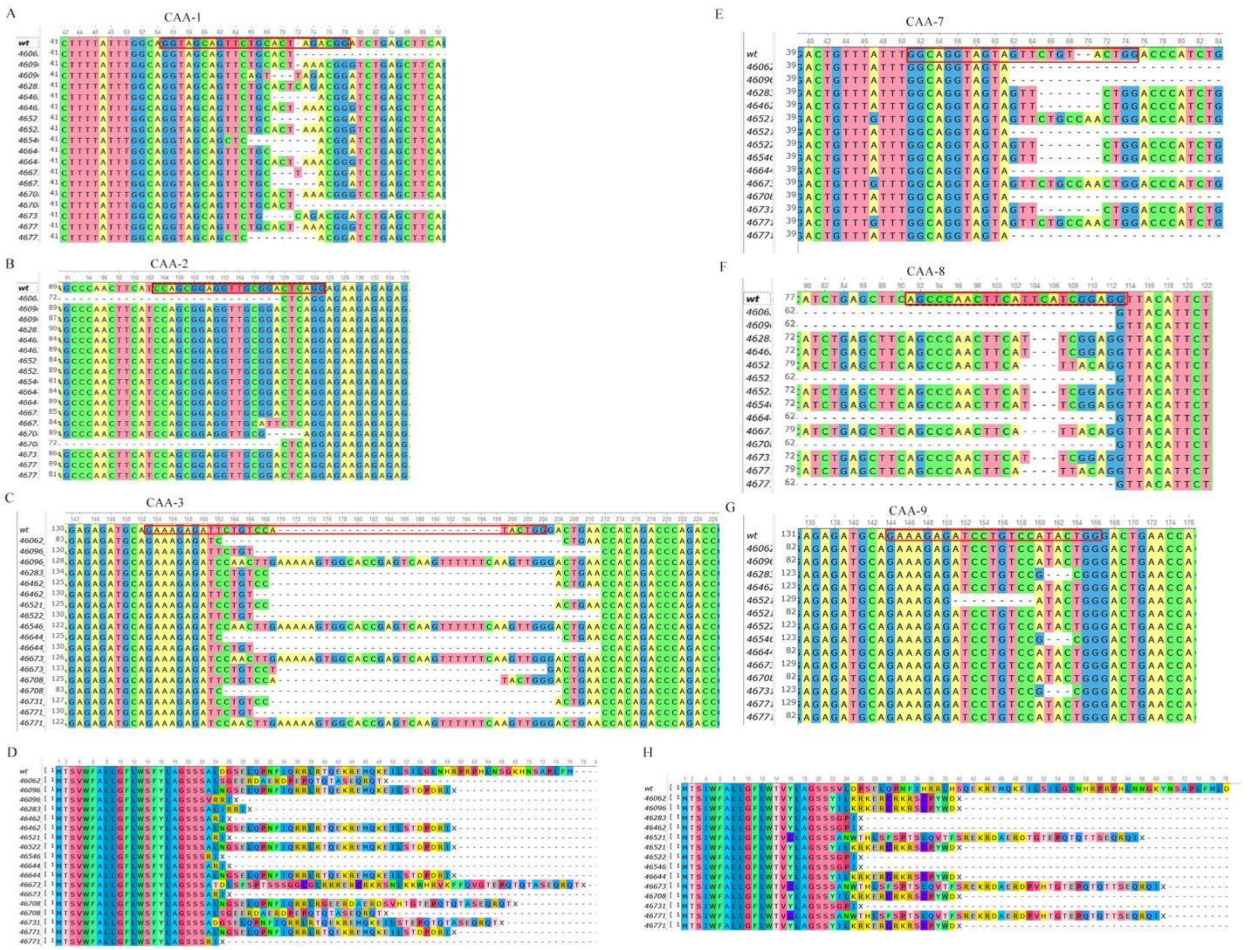

**Supplementary Figure S8. DNA sequences and protein sequences alignments of *bmp6a* exon1 and *bmp6b* exon1 in *bmp6a*$^{-/-}$;*bmp6b*$^{-/-}$ individuals without intermuscular bones in the F$_2$ generation.** CAA-1 (A), CAA-2(B), and CAA-3 (C) were the target sites (marked with red box) for *bmp6a* knockout. (D) Alignments of *bmp6a* protein sequences inferred with DNA sequences in F$_2$ mutants without IMBs. CAA-7(E), CAA-8(F), and CAA-9 (G) were the target sites (marked with red box) for *bmp6b* knockout. (H) Alignments of *bmp6b* protein sequences inferred with DNA sequences in F$_2$ mutants without IMBs. WT represents wild-type, and the others represent mutants.

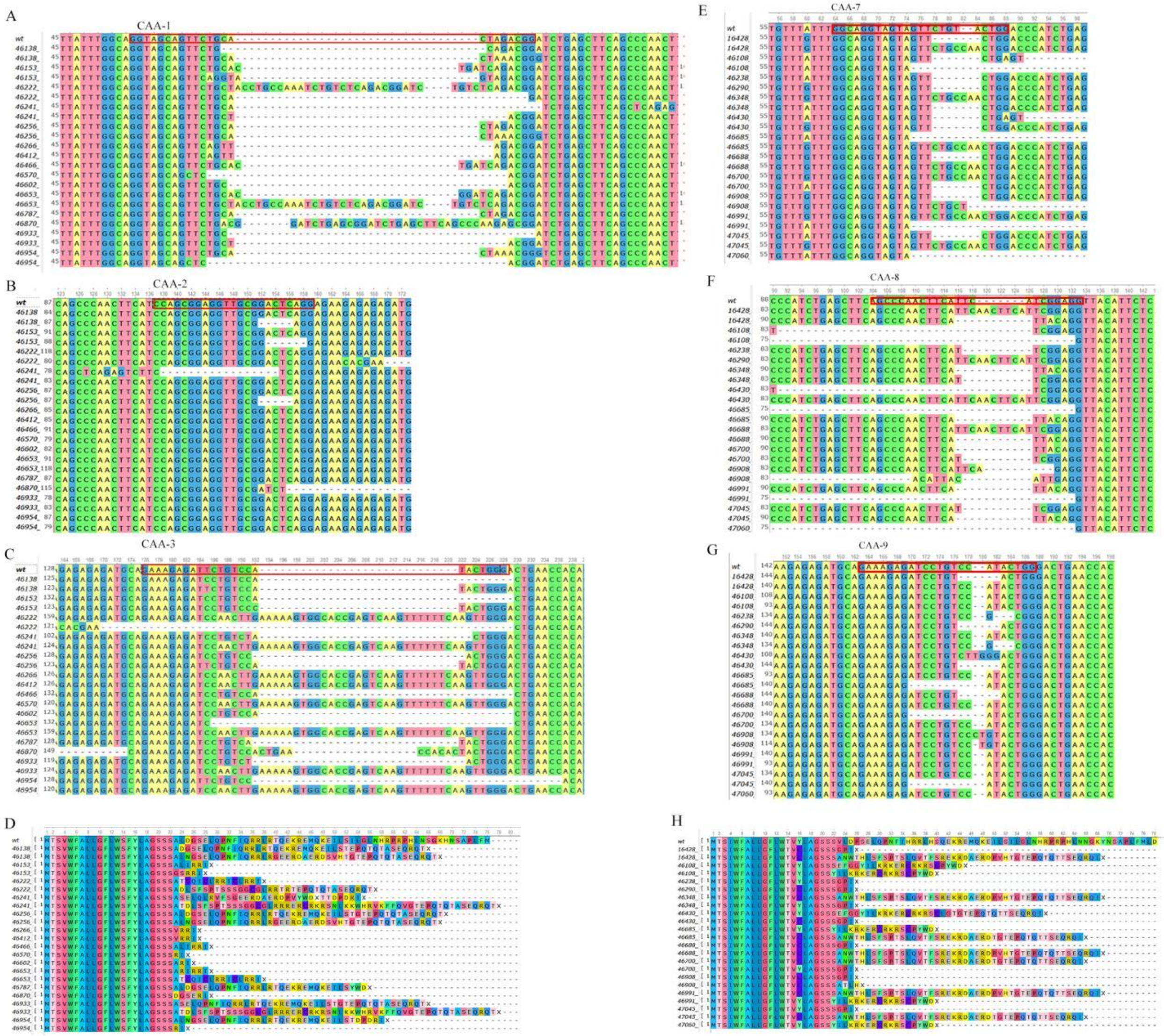

**Supplementary Figure S9. DNA sequence and protein sequences alignments of *bmp6a* exon1 in *bmp6a*$^{-/-}$;*bmp6b*$^{+/+}$ individuals(A-D) and *bmp6b* exon1 in *bmp6a*$^{+/+}$;*bmp6b*$^{-/-}$ individuals(E-H).** CAA-1(A), CAA-2(B), and CAA-3(C) were the target sites (marked with red box) for *bmp6a* knockout. (D) Alignments of *bmp6a* protein sequences inferred with DNA sequences in *bmp6a*$^{-/-}$;*bmp6b*$^{+/+}$ mutants with partial deletion of intermuscular bones. CAA-7(E), CAA-8(F), and CAA-9(G) were the target sites (marked with red box) for *bmp6b* knockout. (H) Alignments of *bmp6b* protein sequences inferred with DNA sequences in *bmp6a*$^{+/+}$;*bmp6b*$^{-/-}$ mutants with partial deletion of intermuscular bones. WT represents wild-type, and the others represent mutants. The phenotype of *bmp6a*$^{-/-}$;*bmp6b*$^{+/+}$ was shown in the top right of Fig 3g, and the phenotype of *bmp6a*$^{+/+}$;*bmp6b*$^{-/-}$ was shown in the bottom left of Fig 3g.

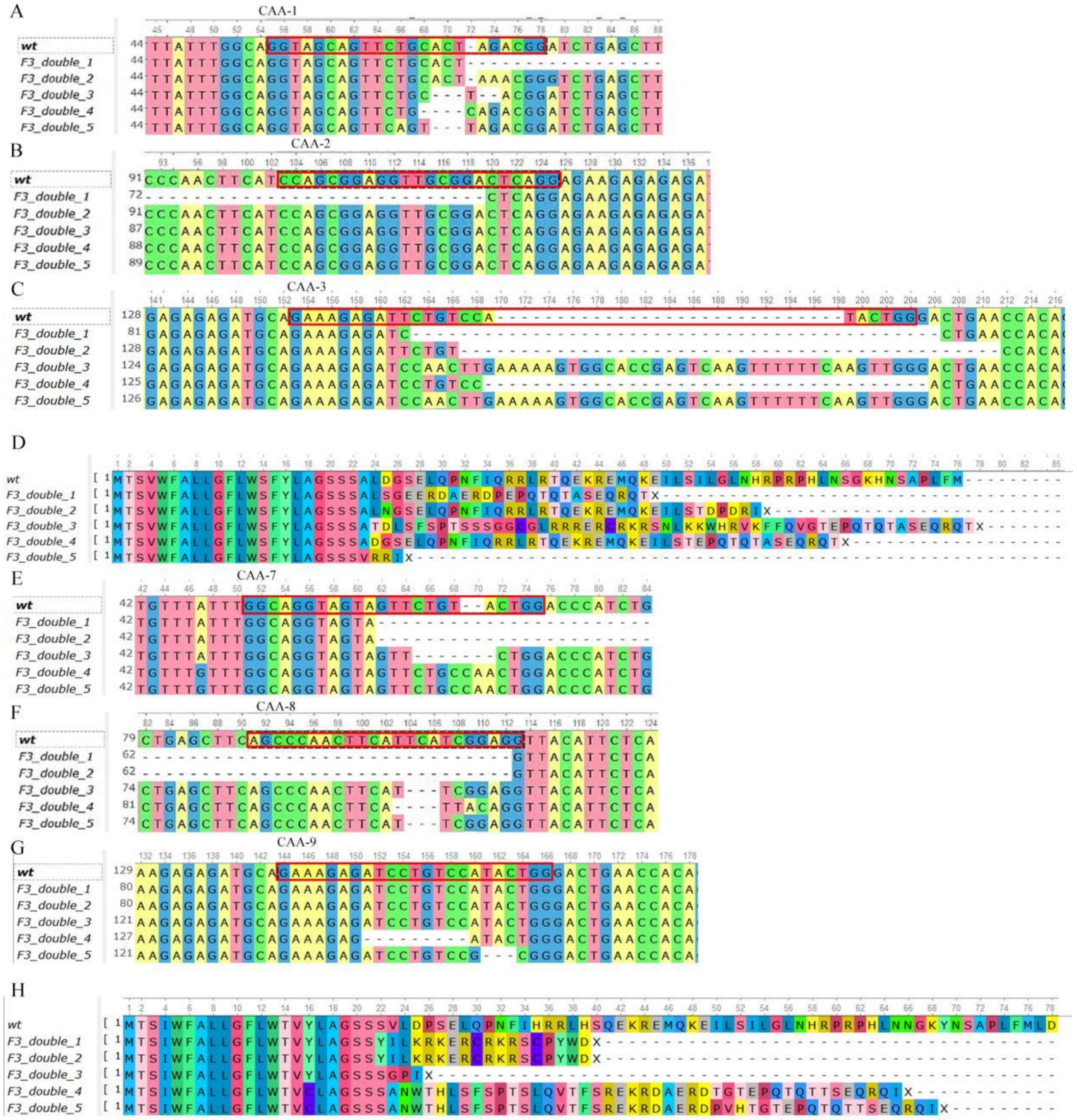


**Supplementary Figure S10. DNA Sequences alignments and protein sequences of *bmp6a* exon1 and *bmp6b* exon1 in *bmp6a*$^{-/-}$;*bmp6b*$^{-/-}$ individuals of the $F_3$ generation.** CAA-1, CAA-2, and CAA-3 were the target sites (marked with red box) for *bmp6a* knockout. CAA-7, CAA-8, and CAA-9 were the target sites (marked with red box) for *bmp6b* knockout. WT represents wild-type, and the others represent mutants.

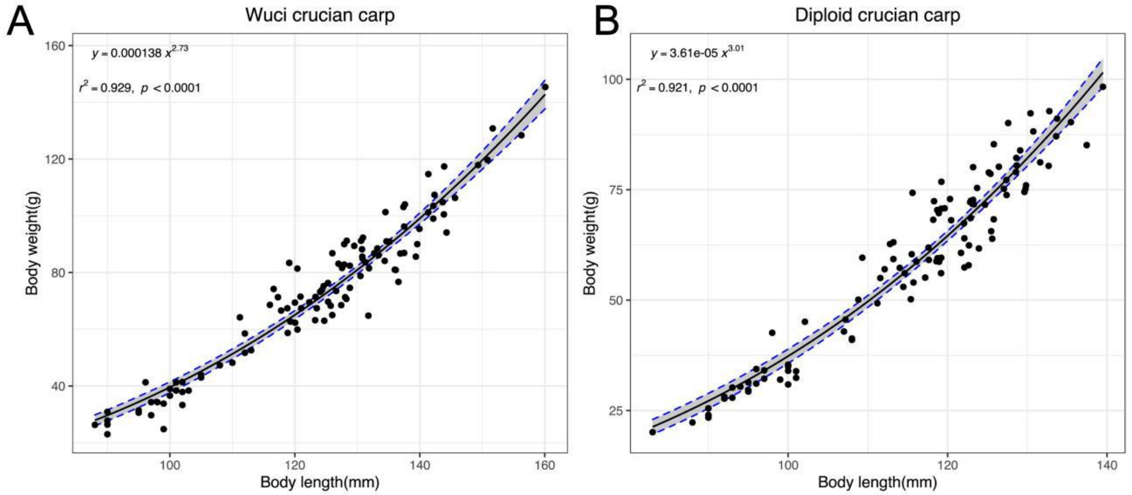


**Supplementary Figure S11. The relationship between body weight and body length.** A: the new strain without intermuscular bones; B: the wild-type crucian carp.

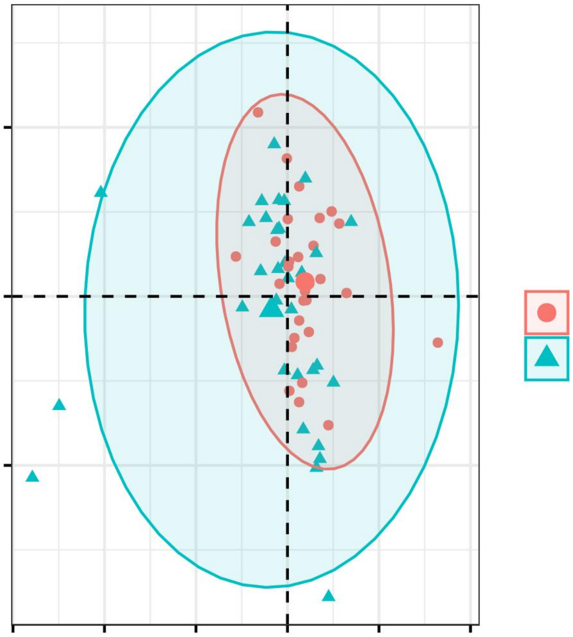

**Supplementary Figure S12. PCA analysis of muscle texture.** WT: wild-type crucian carp (WT), WUCI: the new strain without intermuscular bones.

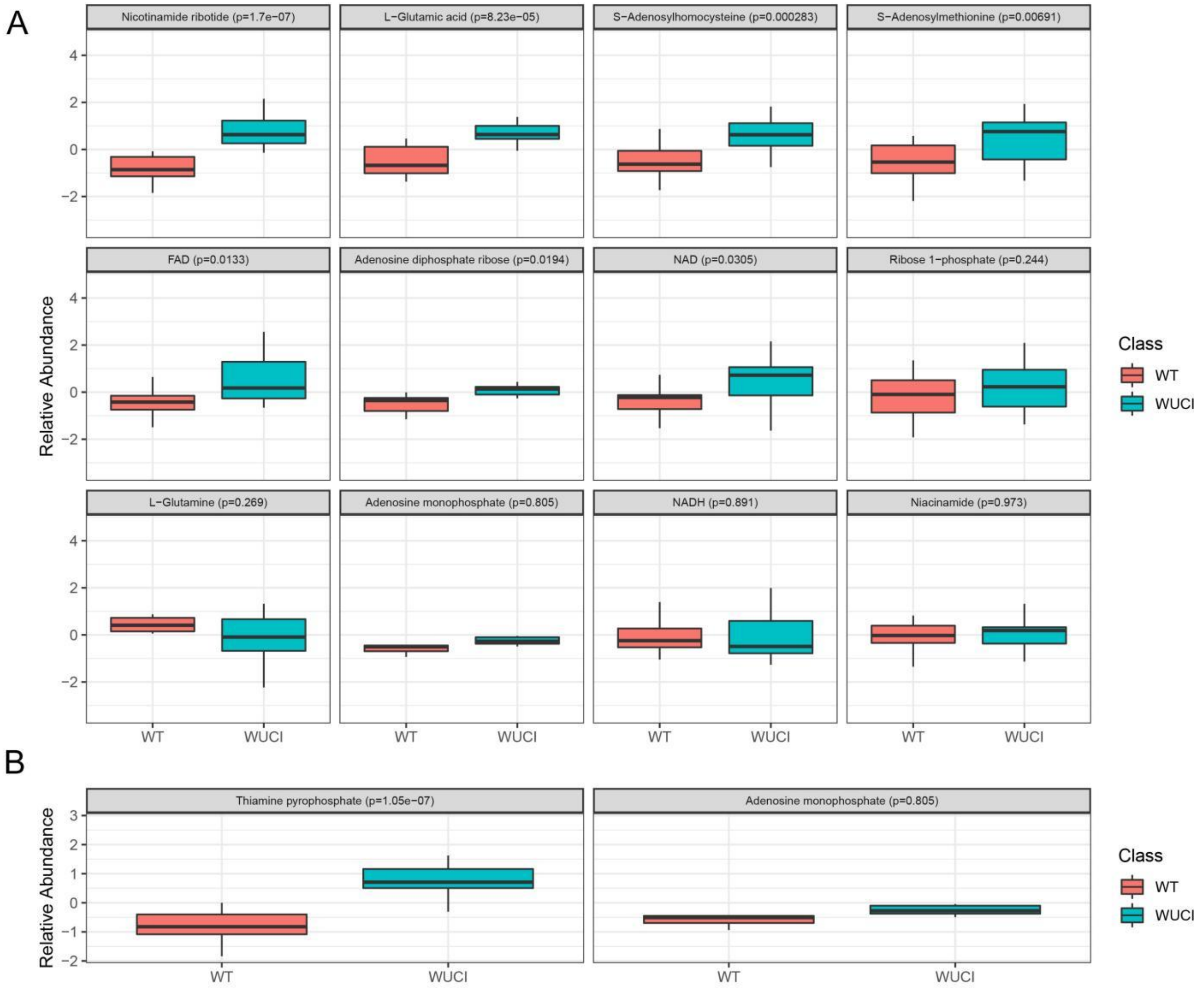


**Supplementary Figure S13. Boxplot of metabolites in Nicotinate and Nicotinamide Metabolism(A) and Thiamine metabolism(B)**

**Supplementary Table S1-S17 was included in a separate Excel file.**
